# Posttraumatic reexperiencing and alcohol use: mediofrontal theta as a neural mechanism for negative reinforcement

**DOI:** 10.1101/2023.07.12.547253

**Authors:** Eric Rawls, Craig A. Marquardt, Spencer T. Fix, Edward Bernat, Scott R. Sponheim

## Abstract

**Objective:** Over half of US military veterans with posttraumatic stress disorder (PTSD) use alcohol heavily, potentially to cope with their symptoms. This study investigated the neural underpinnings of PTSD symptoms and heavy drinking in veterans. We focused on brain responses to salient outcomes within predictive coding theory. This framework suggests the brain generates prediction errors (PEs) when outcomes deviate from expectations. Alcohol use might provide negative reinforcement by reducing the salience of negatively-valenced PEs and dampening experiences like loss.

**Methods:** We analyzed electroencephalography (EEG) responses to unpredictable gain/loss feedback in veterans of Operations Enduring and Iraqi Freedom. We used time-frequency principal components analysis of event-related potentials to isolate neural responses indicative of PEs, identifying mediofrontal theta linked to losses (feedback-related negativity, FRN) and central delta associated with gains (reward positivity, RewP).

**Results:** Intrusive reexperiencing symptoms of PTSD were associated with intensified mediofrontal theta signaling during losses, suggesting heightened negative PE sensitivity. Conversely, increased hazardous alcohol use was associated with reduced theta responses, implying a dampening of these negative PEs. The separate delta-RewP component showed associations with alcohol use but not PTSD symptoms.

**Conclusions:** Findings suggest a common neural component of PTSD and hazardous alcohol use involving altered PE processing. We suggest that reexperiencing enhances the intensity of salient negative PEs, while chronic alcohol use may reduce their intensity, thereby providing negative reinforcement by muting emotional disruption from reexperienced trauma. Modifying the mediofrontal theta response could address the intertwined nature of PTSD symptoms and alcohol use, providing new avenues for treatment.

## 1. Introduction

Combat veterans frequently encounter mental health issues like posttraumatic stress disorder (PTSD) and heavy alcohol use. About 23% of combat veterans have PTSD (Fulton et al., 2015), while 10.5% have alcohol dependence (AD) (Seal et al., 2011). A substantial intersection exists between PTSD and heavy drinking. 50-76% of PTSD-diagnosed veterans fulfill AD criteria (Jakupcak et al., 2010; Panza et al., 2021; Wisco et al., 2014), and it is possible that a larger proportion engage in heavy drinking as a coping mechanism. As such, negative reinforcement (psychological benefit due to avoidance or escape from aversive stimuli or states) (Koob, 2013) likely plays a prominent role in the link between heavy drinking and PTSD. Despite high rates of alcohol use in veterans with PTSD, no studies that we are aware of have identified neural activity related to PTSD and alcohol use that could explain their covariation in military veterans. This study aims to elucidate the neural correlates of posttraumatic symptomatology and heavy drinking by focusing on how combat veterans experience and respond to losses and rewards (i.e., salient stimuli).

Individuals with PTSD perceive their surroundings as more threatening and show attentional biases toward threat (Clauss et al., 2022). Enhanced salience of cues for potential losses and gains is linked to PTSD symptomatology and brain salience and reward system activation (Jia et al., 2023). This investigation, informed by predictive coding (Friston & Kiebel, 2009), examines how PTSD and heavy alcohol use affect brain processing of gains and losses (Kube et al., 2020; Putica et al., 2022). Predictive coding suggests the brain forms predictions (‘priors’) and adjusts them based on deviations from expectations (‘prediction errors;’ PEs). PEs indicate whether outcomes are better (positive PE) or worse (negative PE) than predicted. Traumatic experiences can lead to strong priors about potential threats, intensifying processing of benign stimuli as overly salient and negative. This is linked to intrusive reexperiencing of traumatic events (Kube et al., 2020; Putica et al., 2022), where benign stimuli trigger strong threat representations tied to past experiences, essentially turning them into negative PEs. We suggest alcohol use might reduce the salience of these negative PEs, offering relief from reexperiencing symptoms but risking reinforcing maladaptive drinking behaviors (Berenz et al., 2021; Weiss et al., 2021). Essentially, alcohol’s negatively reinforcing effects (Koob, 2013) may stem from reducing brain responses to negative PEs.

We assessed brain responses to unpredictable gain/loss feedback using electroencephalography (EEG). The EEG shows a mediofrontal feedback-related negativity (FRN), pronounced following loss and appearing 250-350 ms post-feedback (Gehring & Willoughby, 2002). Sometimes FRN measurement overlaps with a similarly timed Reward Positivity (RewP) (Proudfit, 2015). We applied principal components analysis (PCA), a dimension reduction technique, to distinguish the frequency-specific content of ERPs. In the time-frequency domain, the FRN corresponds with theta-band (4-8 Hz) activity and likely reflects the output of the brain’s salience network (Seeley et al., 2007), notably anterior cingulate cortex (Cavanagh & Shackman, 2015). The ACC might enact predictive coding by computing negatively-biased surprise signals (or PEs) that assist with learning (Alexander & Brown, 2019). The theta-FRN, linked to ACC predictive coding mechanisms and indicative of negative emotion and cognitive control, could illuminate how PTSD and heavy drinking influence brain salience processing.

Feedback-locked ERPs also show a RewP, more pronounced for gains than losses (Proudfit, 2015). The RewP responds primarily to positive PEs and contains delta-band (0.5-3 Hz) activity (Cavanagh, 2015). PCA identifies the RewP as a positive component separate from the FRN (Hager et al., 2022; Yin et al., 2018). There is some dissociation between the stimulus-locked P300 and the RewP, as the RewP has a more anterior scalp topography (maximal at Cz) and earlier onset (∼200 ms) than the stimulus P300 (maximal at Pz, onset ∼300 ms). The RewP is nevertheless morphologically and functionally similar to the stimulus-locked P300, which is associated with externalizing personality traits (Bernat et al., 2011) including impulsivity and aggressiveness (Krueger et al., 2005). The P300 has a strong genetic basis reflecting predisposition toward substance use (Benegal et al., 1995; Iacono et al., 2003; Polich & Bloom, 1999). This relationship with externalizing appears also to extend to the RewP (Bernat et al., 2011), underscoring its close relationship to the P300. In this study, RewP/P300 might reflect a neural predisposition for alcohol use rather than the emotional distress associated with PTSD.

Previous studies have shown that AD corresponds with diminished FRN and RewP (Kamarajan et al., 2010), whereas PTSD symptomatology is associated with amplified RewPs (Lieberman et al., 2017). The interplay between PTSD and heavy drinking, and specifically brain responses to salient loss and reward, remains largely uncharted. This study, employing a gambling task, examines gain/loss outcome processing in relation to PTSD and heavy drinking. We focus on theta-FRN and delta-RewP, because they are linked to loss and gain processing. Our post-deployment veteran sample, characterized by prevalent posttraumatic reexperiencing symptoms, offers insights into how emotional dysregulation following trauma and heavy drinking are tied to brain responses to salient stimuli. We hypothesize that the severity of reexperiencing symptoms and heavy drinking will be independently associated with neural salience processing patterns. These results would deepen our understanding of PTSD’s neural underpinnings and suggest a model where heavy drinking maladaptively mitigates the exaggerated salience signaling typical of intrusive reexperiencing.

## 2 Methods & Materials

### 2.1 Participants

The sample consisted of 128 US military veterans who had been deployed to Operations Iraqi Freedom or Enduring Freedom (see *Table 1* for demographics). Recruitment targeted veterans with likely posttraumatic stress disorder (PTSD) diagnoses as well as non-treatment-seeking veterans with similar deployment experiences [see (Davenport et al., 2014) for complete recruitment information]. Study procedures were approved by the Institutional Review Boards at the Minneapolis Veterans Affairs Health Care System and the University of Minnesota, and study participants completed a written informed consent process prior to undergoing the study procedures. No prior publications have involved the EEG data collected using the gambling paradigm that is the focus of this manuscript.

**Table 1.** Demographic and clinical characteristics of sample. Note that demographics are shown split by four groups in order to provide full clinical information on the sample, but primary analyses used continuous severity measures instead of diagnosis-based groups.

| Variable | No PTSD |  |  |  |  |  | PTSD+Subthreshold |  |  |  |  |  |
| --- | --- | --- | --- | --- | --- | --- | --- | --- | --- | --- | --- | --- |
|  | No AD |  |  | AD |  |  | No AD |  |  | AD |  |  |
|  | n | M | SD | n | M | SD | n | M | SD | n | M | SD |
| Total Count | 59 |  |  | 12 |  |  | 38 |  |  | 19 |  |  |
| Female | 8 |  |  | 2 |  |  | 0 |  |  | 0 |  |  |
| Race |  |  |  |  |  |  |  |  |  |  |  |  |
| White | 52 |  |  | 12 |  |  | 35 |  |  | 19 |  |  |
| Black | 2 |  |  | 0 |  |  | 0 |  |  | 0 |  |  |
| Asian | 1 |  |  | 0 |  |  | 0 |  |  | 0 |  |  |
| Multiracial | 4 |  |  | 0 |  |  | 3 |  |  | 0 |  |  |
| Age (years) |  | 33.42 | 8.22 |  | 30.83 | 7.57 |  | 31.16 | 8.26 |  | 31.42 | 7.30 |
| Education (years) |  | 5.44 | 0.70 |  | 4.83 | 0.71 |  | 5.21 | 0.66 |  | 5.21 | 0.79 |
| Depressive Disorder Diagnosis | 5 |  |  | 3 |  |  | 16 |  |  | 9 |  |  |
| mTBI Experienced | 19 |  |  | 6 |  |  | 21 |  |  | 12 |  |  |
| CAPS Intrusive Reexperiencing |  | 10.22 | 4.62 |  | 9.14 | 4.53 |  | 16.76 | 5.79 |  | 19.37 | 6.29 |
| CAPS Avoidance |  | 3.83 | 2.85 |  | 4.14 | 3.85 |  | 8.68 | 3.11 |  | 9.37 | 3.73 |
| CAPS Dysphoria |  | 12.27 | 7.31 |  | 9.43 | 3.60 |  | 26.13 | 8.91 |  | 29.37 | 8.64 |
| CAPS Hyperarousal |  | 4.89 | 3.45 |  | 7.43 | 3.64 |  | 7.92 | 3.40 |  | 9.11 | 2.69 |
| AUDIT-C |  | 4.14 | 2.10 |  | 8.25 | 1.54 |  | 4.00 | 2.61 |  | 6.89 | 2.18 |
| Above AUDIT-C Cutoff | 36 |  |  | 12 |  |  | 17 |  |  | 17 |  |  |
| MN-BEST Blast mTBI Severity |  | 1.00 | 1.71 |  | 0.92 | 1.16 |  | 2.03 | 3.00 |  | 1.89 | 2.16 |

PTSD = posttraumatic stress disorder, AD = alcohol dependence, mTBI = mild traumatic brain injury, N = count, M = mean, SD = standard deviation, CAPS = Clinician-Administered PTSD Scale, AUDIT-C = Alcohol Use Disorders Identification Test, MN-BEST = Minnesota Blast Exposure Screening Tool. “+Subthreshold” reflects individuals who meet criteria for at least one symptom from each symptom domain of DSM-IV PTSD. The AUDIT-C cutoff was $\geq 4$ for men and $\geq 3$ for women.

### 2.2 Clinical Assessment

Trained and supervised interviewers conducted assessments for psychopathology using the Structured Clinical Interview for DSM-IV Axis I Disorders [SCID-I; (First & Gibbon, 2004)]. Interviewers characterized posttraumatic stress symptoms using the Clinician-Administered PTSD Scale for DSM-IV [CAPS, fourth edition; (Blake et al., 1995; Weathers et al., 2001)]. We subdivided the CAPS into four subscales based on previous meta-analytic research on the factor structure of the CAPS (Palmieri et al., 2007; Simms et al., 2002; Yufik & Simms, 2010), which provided measures of the severity of intrusive reexperiencing (B1 - B5), avoidance (C1, C2), dysphoria (C3 - D3), and hyperarousal symptoms (D4, D5). Participants only completed the full CAPS if they met criteria A1/A2 and B of the CAPS using DSM-IV-TR criteria (i.e. endorsed a traumatic event with an intense emotional response and later experienced intrusive reexperiencing); as such, dimensional analyses included a subsample of 82 subjects who reported a traumatic event with current reexperiencing (Marquardt et al., 2021).

Consensus teams, including at least one licensed doctoral-level clinical psychologist, reviewed all available research and clinical information to generate consensus diagnoses which included PTSD, subthreshold PTSD, and alcohol dependence (AD). Individuals were given a subthreshold PTSD designation if they endorsed at least one symptom in each DSM-IV-TR symptom grouping for PTSD, consistent with rating schemes meant to increase sensitivity for clinically meaningful presentations of PTSD symptoms (Marquardt et al., 2022). We assessed the severity of alcohol use with the Alcohol Use Disorders Identification Test (AUDIT)-C (Saunders et al., 1993), a 3-item self-report measure of frequency of alcohol use, amount of alcohol use, and frequency of binge drinking. The scale has a maximum score of 12, and the cutoff for clinically meaningful drinking is a score of 4 for men or a score of 3 for women. We assessed for a history of mild traumatic brain injury (mTBI) using the semi-structured Minnesota Blast Exposure Screening Tool [MN-BEST] (Nelson et al., 2011), focusing on the three most severe self-identified deployment-related blast exposure events. We achieved consensus on mTBI via assessment teams that included at least one licensed clinical neuropsychologist. Importantly, the study recruitment criteria used a diagnosis of pre-deployment psychopathology as part of exclusion criteria, thus the clinical presentations of psychopathology assessed in the present study are likely to have been acquired post-deployment [see (Davenport et al., 2014)].

### 2.3 Gambling Task

Participants completed a gambling paradigm originally described in (Gehring & Willoughby, 2002). Each trial offered participants a two-option forced choice. Options were 5 or 25 cents, and could be paired in any fashion (i.e. 5/5, 5/25, or 25/25) with all pairs being equiprobable. Choices were presented within black squares which remained on the screen until participants selected one option. One hundred ms following the choice, each square turned red or green (*Figure 2A*). If the chosen option turned green, the indicated amount was added to the participant’s running score. If the chosen option turned red, the indicated amount was instead subtracted from the participant’s running score. The color of the unchosen option also changed, to indicate what the outcome would have been if the participant had instead chosen that option. Participants completed 256 trials, divided into 8 blocks with self-paced breaks in between.

This task required approximately 20 minutes to complete. Participants received additional monetary compensation at the end of the study session equivalent to their total United States dollar amount earned during this task. An important feature of the task was the unpredictable nature of choice feedback. The primary behavioral outcome was risky choice proportion, defined as the percentage of times a participant chose the ‘25’ option when presented with a choice between ‘5’ and ‘25.’ This risky choice proportion was calculated separately for trials following gains and losses Participants are often more risk prone following losses compared with gains (Gehring & Willoughby, 2002).

### 2.4 EEG Acquisition, Preprocessing, and Time-Frequency PCA Analysis

EEG was sampled at 1024 Hz using a 128-channel BioSemi ActiveTwo EEG system, acquired reference-free (via CMS/DRL sensors). EEG data were preprocessed and analyzed exactly as described in (Bernat et al., 2011); the following is paraphrased. EEG were imported and re-referenced to linked mastoids, epoched surrounding gain/loss feedback [−1,000 to 2,000 ms; extended to mitigate edge artifacts (Cohen, 2014)], and baseline corrected (-150 - 0 ms). Disconnected sensors were identified and interpolated. Ocular artifacts were removed via regression (Gratton et al., 1983). Remaining artifacts were removed by deleting trials where frontal activity (sensors C12/C25) exceeded 100 μV within a 1,500-ms poststimulus window or an 800-ms prestimulus window. Additional movement and other artifacts were identified and removed via visual inspection. We then calculated ERPs at each sensor separately for gain/loss trials.

We reduced ERP dimensionality using time-frequency principal components analysis [tf-PCA; (Bernat et al., 2005; Buzzell et al., 2022)] calculated using the Psychophysiology Toolbox (PTB; http://www.ccnlab.umd.edu/Psychophysiology_Toolbox/). To allow tf-PCA to define the boundary between delta and theta, we pre-filtered ERP waveforms using a 4-Hz low-pass Butterworth filter for delta, and 2-Hz high-pass Butterworth filter for theta (third order, zero-phase). Filtered waveforms were transformed to a TF representation using the binomial reduced interference distribution (Jeong & Williams, 1992). We vectorized TF surfaces into a matrix of dimensions subjects-by-TF points and applied PCA to the covariance matrix. We chose the number of components to retain using an eigenvalue scree plot, retaining one delta component (62% of variance) and three theta components (22%, 21%, and 9% of variance). We applied a varimax rotation (Bernat et al., 2005, 2011) to the loadings then reshaped them into TF matrices. Delta loadings mapped well onto the scalp distribution and timing of the central RewP, and the second theta-band component mapped well onto the scalp distribution and timing of the FRN. The remaining theta components were not analyzed as they reflected the occipital N1 ERP component and a 2.5-3 Hz non-FRN oscillation. Dependent theta-FRN and delta-RewP values were calculated by averaging PC-weighted TF surfaces at sensors where component activation were maximal (Cz for delta, FCz for theta).

### 2.5 Statistical Analysis

Statistics were conducted in R version 4.2.3. We had three outcome measures: risky choices, delta-RewP, and theta-FRN activation. Our sample had a wide age range (22 - 59 years old), so we screened DVs for associations with age. Theta-FRN was associated with age (*r* = -.25, *p* < .001), so theta-FRN analyses contrived for age. We used robust linear mixed-effects models (rLMMs) fit with the ‘robustlmm’ package, version 3.0-4 (Koller, 2016) because theta-FRN and delta-RewP were highly skewed (skewness = 1.9 and 1.7 respectively) relative to the assumptions of non-robust LMMs (Arnau et al., 2013). We estimated rLMM *p*-values using robust *t*-statistics and Kenward-Roger approximated degrees-of-freedom.

RLMMs analyzing delta-RewP and theta-FRN had a within-subject factor of Outcome (gain/lose). RLMMs analyzing risky choice percentage had a within-subject factor of Previous Outcome (previous gain/ previous loss). RLMMs testing brain-behavior associations had a within-subject factor of Previous Outcome (previous gain/ previous loss), and included delta-RewP and theta-FRN as continuous predictors.

RLMMs also included between-subjects factors describing clinical presentation. In the first analysis, we simultaneously entered between-subjects factors for clinical diagnoses of PTSD, mTBI, and AD. In the second analysis, we simultaneously entered continuous between-subjects variables consisting of the four CAPS subscales (intrusion/ avoidance/ dysphoria/ hyperarousal), AUDIT-C, and blast mTBI severity. Noting that individual CAPS subscales are associated with each other, we assessed for multicollinearity using variance inflation factor (VIFs) calculated for each model using the ‘performance’ package version 0.10.8 (Lüdecke et al., 2021). All VIF were < 2.5, with a criterion of VIF ≥ 5 considered evidence of multicollinearity.

All IVs and DVs were *z*-scored to reduce multicollinearity and obtain standardized effect size estimates. All models contained a random intercept per participant and interaction terms between the within-subjects Outcome factor and all between-subjects factors, but did not include interactions of between-subjects factors. Post-hoc characterization of significant interactions used the ‘emmeans’ package, version 1.7.4-1 (Lenth et al., 2022).

## 3. Results

### 3.1 Risky Gambling Behavior is Related to Alcohol Use

A diagram of the gambling task and of risky choice rates is shown in *Figure 1*. Risky choice behavior on the gambling task showed an expected main effect of Outcome (gain/loss) in all analyses, *ts* ≥ 7.13, *ps* < .001, indicating higher risky choice behaviors following loss outcomes. Group analyses focusing on Diagnosis (yes/no, PTSD/mTBI/AD) showed no effects of PTSD or mTBI, but revealed a main effect of an AD diagnosis, *t*(124) = 2.34, *p* = .021, indicating overall higher risky choice behavior in participants with AD. Likewise, a dimensional analysis focusing on symptom severity (CAPS subscales, mTBI severity, AUDIT-C score) revealed a main effect of AUDIT-C, *t*(75) = 2.03, *p* = .046, indicating overall higher risky choice behavior in participants with greater alcohol consumption. This analysis failed to show any independent effects of PTSD symptomatology or mTBI severity on risky choice behaviors within the same models.

**Figure 1.**
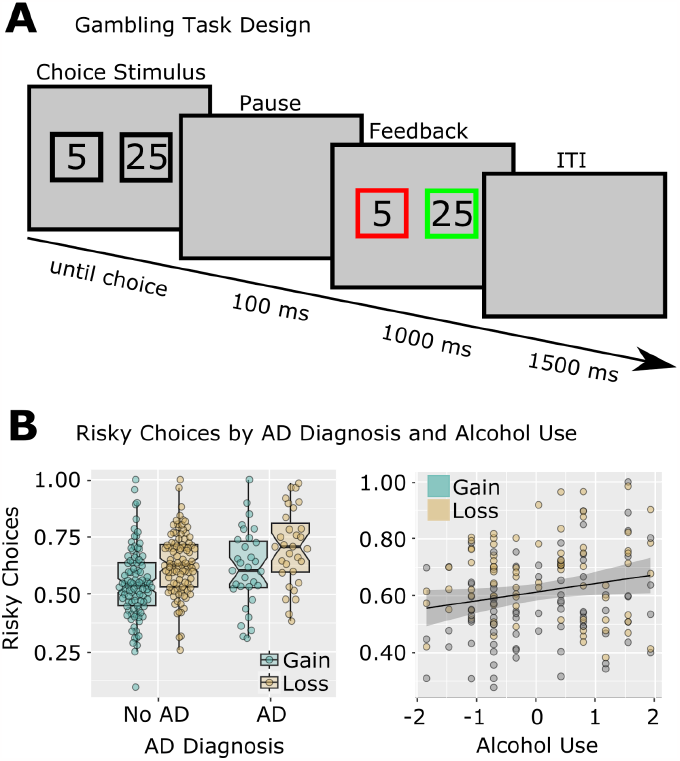
Risky Gambling Behavior is Related to Alcohol Use in Previously Deployed Veterans. A: Design of the modified gambling task. B: Risky choices were increased following losses compared to gains. Individuals with AD and with higher AUDIT-C scores made more risky choices. Note that individual data points are shown to differentiate gain/loss observations, but all statistics were main effects over both Gain/Loss conditions (thus there is only one regression line, rather than separate regressions for gain and loss). AUDIT-C was standardized for analysis and plotting; risky choice proportions were standardized for analysis but not for plotting.

**Figure 2.**
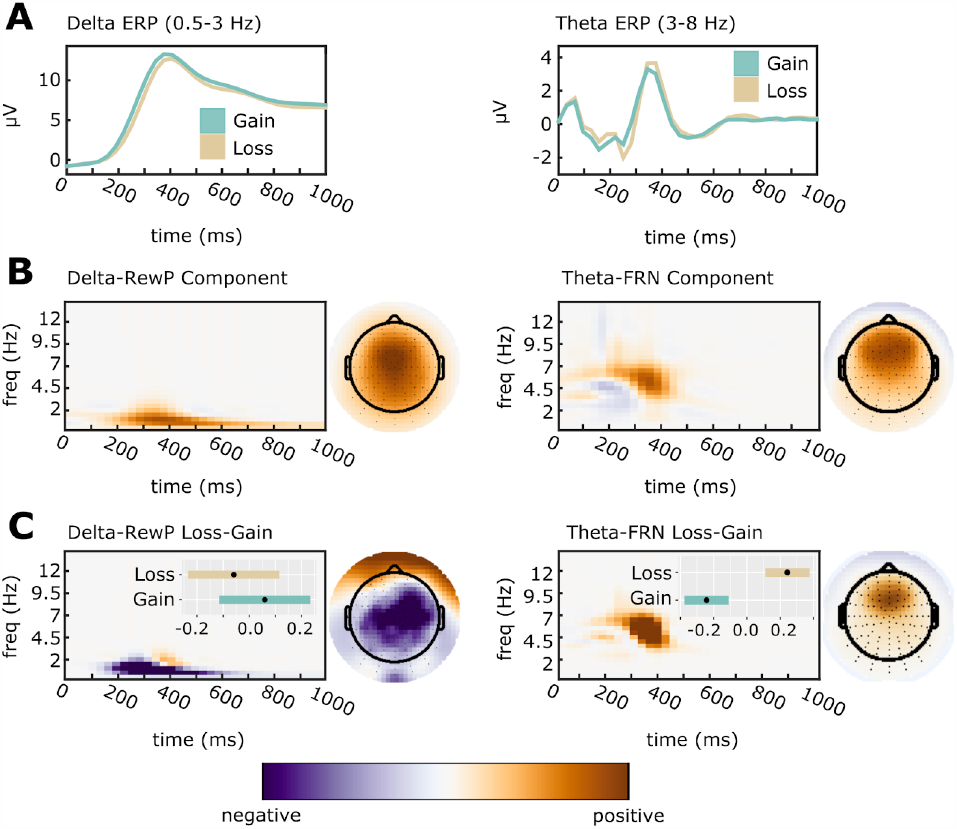
Time-Frequency Principal Components Analysis was applied to separate overlapping ERP activation. All TF surfaces and topoplots are plotted with zero (white) as midpoint. Data units are arbitrary since plots depict PC-weighted power; thus, each plot is scaled to the range of the data. A: Averaged ERP waveforms were filtered into delta (0.5-3 Hz; Cz electrode) and theta (4-8 Hz; FCz electrode) bands. B: ERP waveforms were decomposed, and components reflecting the delta-RewP and theta-FRN response were selected for further analysis based on their PC weights. Components were selected for analysis based on an average over gain/loss conditions. C: To confirm the selected components, we calculated topographic maps and time-frequency surfaces for the average subtraction of loss-gain loadings. As expected, delta-RewP showed greater activation for gains than for losses (left panel), while theta-FRN showed greater activation for losses than for gains (right panel). Inset bars indicate estimated marginal means (EMMs) and associated standard errors for component averages. EMMs are for z-scored component amplitudes fit with a random effects model that accounts for subject-specific intercepts.

### 3.2 Delta-RewP is Related to Amount of Alcohol Use

The tf-PCA separation of delta-band RewP from theta-band FRN is shown in *Figure 2*. Our analysis of time-frequency delta PC-weighted activation (i.e. the centro-parietal delta-band activity underlying the RewP) demonstrated a main effect of Outcome for all analyses, *t*s ≥ 3.53, *p*s ≤ .002, indicating relatively greater activation for gains compared to losses. Group analyses focusing on Diagnosis (yes/no, PTSD/mTBI/AD) showed no results. A dimensional analysis focusing on symptom severity (CAPS subscales, mTBI severity, AUDIT-C score) revealed a significant main effect of AUDIT-C total score, *t*(75) = -2.01, *p* = .048, indicating decreasing delta-RewP activation with increasing hazardous drinking, standardized AUDIT-C fixed effect estimate = -.19, 95% CI = [-.381, -.001]. There were no effects of continuous measures of PTSD or blast-related mTBI severity. Thus, this analysis revealed that blunted delta-RewP activation was related to increases in hazardous drinking, but was unrelated to PTSD or mTBI (*Figure 3A*).

**Figure 3.**
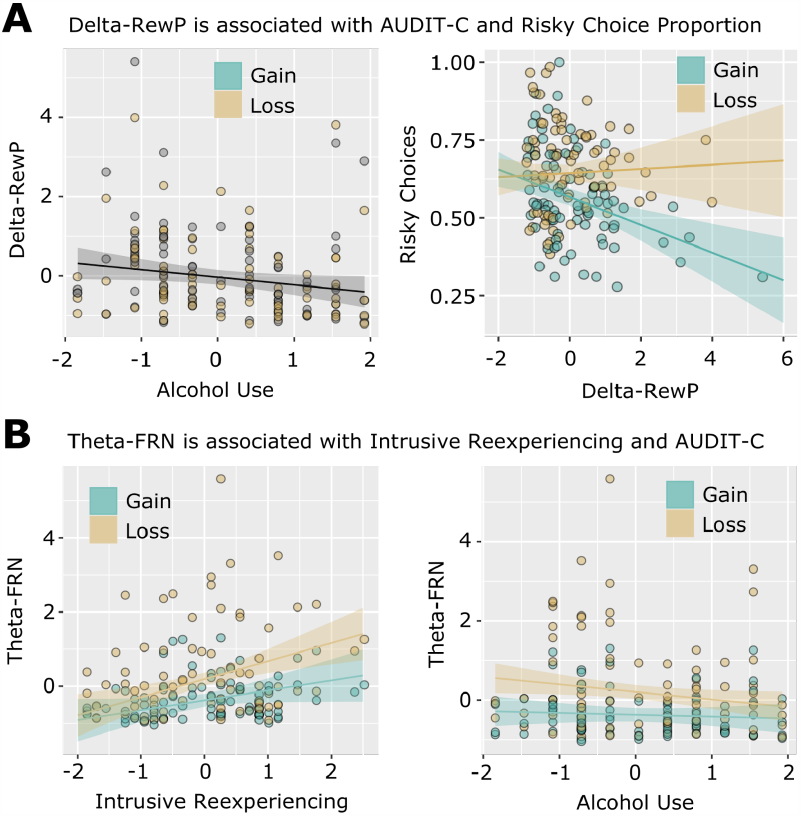
Delta and theta feedback components are related to alcohol use, intrusive reexperiencing, and risky choices in combat veterans. A: Delta-RewP activation was negatively associated with AUDIT-C scores and with risky choices following gains. Delta-RewP and Alcohol Use (AUDIT-C) were standardized for analysis and plotting; risky choice proportions were standardized for analysis but not for plotting. Note that for the left panel, individual data points are shown to differentiate gain/loss observations, but statistics indicate a main effect over both Gain/Loss conditions (thus there is only one regression line, rather than separate regressions for gain and loss). B: Theta-FRN activation was associated with less severe alcohol use (AUDIT–C scores), and more Intrusive Reexperiencing symptoms related to traumatic events. Theta-FRN, Intrusive Reexperiencing, and Alcohol Use (AUDIT-C) were standardized for analysis and plotting.

### 3.3 Opposing Effects of Intrusive Reexperiencing and Alcohol Use on Theta FRN

The tf-PCA separation of theta-band FRN from delta-band RewP is shown in *Figure 2*. Our analysis of time-frequency theta PC-weighted activation (i.e. the mediofrontal theta-band activity underlying the FRN) demonstrated a main effect of Outcome for all analyses, *ts* ≤ -8.37, *ps* < .001, indicating greater activation for losses than gains. Group analyses focusing on Diagnosis (yes/no, PTSD/mTBI/AD) showed no results. Our analysis of individual differences using dimensions of PTSD symptoms, alcohol use, and blast-related mTBI yielded a main effect of Intrusive Reexperiencing, *t*(75) = 2.93, *p* = .004. The main effect of Intrusive Reexperiencing was qualified by a significant interaction with Outcome, *t*(75) = -2.09, *p* = .040. Finally, the model also simultaneously identified a significant interaction between AUDIT-C and Outcome, *t*(75) = 2.09, *p* = .040. Post hoc examination revealed that greater Intrusive Reexperiencing severity was associated with enhanced theta activation during loss conditions, standardized fixed-effect estimate = 0.46, 95% CI = [0.20, 0.71], *t*(104) = 3.52, *p* < .001, but not gain conditions, standardized fixed-effect estimate = 0.24, 95% CI = [-0.02, 0.49], *t*(104) = 1.94, *p* = .065 (*Figure 3b)*. Post hoc examination of the significant AUDIT-C-Outcome interaction indicated that more alcohol use was associated with reduced theta activation during loss conditions, standardized AUDIT-C fixed-effect estimate = -0.19, 95% CI = [-0.35, -0.03], *t*(104) = -2.29, *p* = .022, but not gain conditions, standardized AUDIT-C fixed-effect estimate = -0.05, 95% CI = [-0.21, 0.11], *t*(104) = -0.60, *p* = .546 (*Figure 3B*). This analysis revealed no effects of blast-related mTBI severity. As such, loss processing as embodied in frontal midline theta is simultaneously linked in opposing ways to the severity of PTSD-related intrusive reexperiencing (positive association) and elevated hazardous alcohol use (negative association) in previously deployed combat veterans.

### 3.4 Delta-RewP, but not Theta-FRN, is Related to Risky Choice Behavior

As previously noted, risky choice behavior on the gambling task showed an expected main effect of Outcome (gain/loss) in all analyses that indicated higher risky choice behaviors following loss outcomes (that is, loss feedback precipitated increased risky choices on the following trial). We next examined whether these risky choice behaviors following gains and losses were differentially associated with gain-related delta-RewP activation or loss-related theta-FRN activation. We observed a significant interaction between Outcome (Previous gain/Previous Loss) and delta-RewP activation, *t*(129.37) = -4.40, *p* < .001. This was due to a significant negative association between delta-RewP and risky gambles following gains, standardized delta-RewP fixed-effect estimate = -.29, 95% CI = [-.44 -.14], *t*(220) = -3.86, *p* < .001 (*Figure 3a*). There was no association between delta-RewP and risky choices following loss feedback, *p* = .99. Similarly, there was no association between theta-FRN activation and risky choices, *p* > .27. This analysis clarifies that decreased delta-band processing of gains is associated with increased risk-taking behaviors on trials immediately following gains. That is, decreased delta activation is predictive of individual differences in risky decision making. Theta-band processing of losses is not similarly predictive of risk-taking.

## 4 Discussion

In our study of neural responses to gains and losses in US military veterans, we found that the neural processing of loss is differentially associated with dimensional measures of intrusive reexperiencing of trauma and alcohol consumption. These associations were unapparent in the categorical analyses of PTSD and alcohol dependence diagnoses. Intrusive reexperiencing, one of the cardinal symptom domains of PTSD, was associated with enhanced mediofrontal theta loss signaling, indicating increased salience for negative outcomes. Concurrently, increased alcohol use was linked to reduced theta loss signaling, suggesting that heavy drinking may serve as a maladaptive coping mechanism to dampen heightened salience. Decreased delta-band signaling during gains was associated with heavy alcohol use, and was predictive of risky choices following gains on the gambling task. Results support using dimensional measures to parse the heterogeneous clinical presentations of PTSD into elements that align more closely with neural mechanisms of salience processing, potentially offering more precise intervention targets. Similarly, quantifying the degree of alcohol use appears more informative than solely relying on traditional diagnostic categories.

Predictive coding theories suggest that the brain generates future predictions (“priors”) and minimizes prediction error (PE) by updating these estimates using experience (Friston & Kiebel, 2009). In the context of PTSD, negative future predictions may be particularly intense, leading to enhanced processing of negatively valenced information, or in predictive coding terms, elevated signaling of negative prediction errors (Kube et al., 2020; Putica et al., 2022). This heightened sensitivity to negative PEs can be seen in the enhancement of theta-FRN power for loss compared to gain outcomes. In the following, we argue in favor of predictive coding as an explanatory framework for the observed associations between posttraumatic reexperiencing, alcohol use, and theta-FRN signaling.

The ACC is a crucial node in the brain’s salience network (Seeley et al., 2007), and plays a role in cognitive control (Carter, 1998), processing negatively-valenced information (Cavanagh & Shackman, 2015; Shackman et al., 2011), and valuation (Shenhav et al., 2013). The ACC is argued to constrain predictive coding within the frontal cortex by computing surprise signals (or PEs) that assist with learning models of the environment (Alexander & Brown, 2019). These PEs are neither entirely positively or negatively valenced, but are primarily characterized by a deviation from expectations, necessitating updating an internal model (Alexander & Brown, 2019). Mediofrontal event-related potentials in theta frequencies (4-8 Hz) are believed to originate in the ACC (Cavanagh & Shackman, 2015). The theta-band activity underlying the mediofrontal FRN is potentiated by losses compared to wins in simple gain-maximization gambling tasks, but broader analyses suggest the FRN more generally reflects the degree of surprise associated with outcomes (Hager et al., 2022; Hird et al., 2018; Rawls et al., 2020; Talmi et al., 2013).

A primary finding of our work is that enhanced theta-FRN signaling during loss processing is positively associated with the severity of posttraumatic reexperiencing. The relationship between the reexperiencing aspects of PTSD and brain salience signaling can be viewed through various theoretical lenses. Fear extinction models suggest PTSD arises from persistent fear responses that exhibit a tendency to overgeneralize to inappropriate contexts (Duits et al., 2015; Zuj et al., 2016), leading to exaggerated salience responses to everyday stimuli. Attentional control theories (Marquardt et al., 2022; Schoorl et al., 2014) propose that PTSD is linked to a failure regulating attention towards negative stimuli. These theories, along with the predictive coding framework, all predict that reexperiencing should be associated with enhanced brain salience signaling for negatively-valenced information.

Yet, our analysis of alcohol use adds nuance to these perspectives and clarifies existing theoretical frameworks about the neural consequences of heavy alcohol use in the context of emotional distress. It is important to note that the primary variable of interest in these models was reported average alcohol use, rather than acute alcohol intoxication. Fear extinction theories predict long-term drinking should enhance rather than suppress salience responses because chronic drinking impairs extinction (Holmes et al., 2012; Smiley et al., 2021); this is in contrast to a short-term negative reinforcement explanatory model. Similarly, attentional control theories predict long-term drinking should enhance salience responses by disrupting attentional control (Goldstein & Volkow, 2011). Plus, chronic alcohol consumption is associated with increased, rather than decreased, negative emotional reactivity (Goldstein & Volkow, 2011; Zilverstand et al., 2018). Thus, given some of the existing findings on people with alcohol dependence, one might predict that heavy drinking, in individuals with current posttraumatic reexperiencing, should be positively associated with even greater loss salience signaling.

This prediction is inconsistent with the pattern we report. Instead, when modeled simultaneously with PTSD symptom severity, we found that increased drinking was linked to reduced salience signaling. We interpret these effects as evidence that heavy alcohol use is indeed associated with reduced intensity of salient negative PEs. Notably, this effect was not present when alcohol use was modeled separately from PTSD symptoms. This suggests the neural impacts of negative reinforcement drinking in the context of posttraumatic psychopathology might not be noticeable unless covarying for that psychopathology. One potential mechanism underlying this effect could be that alcohol use in the longer term changes the intensity of negatively-biased predictions. If this theorized mechanism were at play, it would imply that alcohol use should be associated with decreased salience signaling during loss, as increasing alcohol consumption would reduce the intensity of negative priors in individuals with PTSD. In line with this interpretation, prior evidence indicates that individuals with AD have lower anticipatory brain activity prior to rewards, suggesting reduced ability to make accurate predictions in these contexts (Luijten et al., 2017).

Our findings also reveal associations between heavy drinking, brain processing indexed by the Reward Positivity [RewP], and risky choices following gains. The delta-band activity underlying the RewP primarily reflects positive PEs (Cavanagh, 2015; Sambrook & Goslin, 2015, 2016), indexing the degree to which rewards exceed expectations. The delta-RewP was inversely correlated with risky choices following gains. This suggests that diminished positive PE signaling could promote risk-seeking behavior. PEs represent violations of expectations, and we intrinsically seek to minimize the magnitude of expectancy violations (PEs) during value-based decision-making (Friston & Kiebel, 2009; Putica et al., 2022). It follows that higher PE signaling should promote less risky decision-making, since in this paradigm, the definition of “risky” rests solely on the magnitude of the choice stimulus (Gehring & Willoughby, 2002). Interestingly, while heavy drinking was associated with reduced delta-RewP signaling, delta-RewP was not associated with PTSD symptom severity. This suggests that the mechanism driving the association between alcohol use and delta-RewP amplitude may not be rooted in a self-medication or negative reinforcement strategy. Instead this might indicate a separate neurally-based impairment important for explaining a broader pattern of diminished response to PEs. Together with the theta-FRN results, heavy alcohol use appears to be associated with reduced neural salience signaling for negative and positive PEs alike via separate mechanisms.

The RewP is distinguished from the P300, a ubiquitous brain potential observed following unexpected or salient stimuli, by its earlier onset and more anterior scalp distribution. However, our delta-band component shows a broad scalp topography and extended timing akin to the P300, raising the possibility that our component contains both RewP and P300 activity. Reduced P300 amplitudes reflect externalizing personality traits (Gilmore et al., 2010; Patrick et al., 2006), including impulsivity, aggressiveness, disinhibition, and risky or antisocial behaviors (Krueger et al., 2005; Patrick & Drislane, 2015). P300 amplitudes also reflect a genetic risk for alcoholism (Benegal et al., 1995; Iacono et al., 2003; Polich & Bloom, 1999). As such, the negative association between delta power and alcohol use could be explained not by reduced positive PE signaling, but instead by previously known genetic and externalizing influences on P300 amplitude. Future investigation, perhaps with alternative methods focusing on separating the RewP from the P300, will be needed to resolve these alternative interpretations.

Despite informative findings, there are limitations to our study. Our cross-sectional sample precludes assessing whether theta-FRN associations are a consequence of, or risk/vulnerability factor for, posttraumatic stress (Bonanno, 2005; Luthar et al., 2000; Polusny et al., 2017). Future longitudinal studies involving new military recruits before and after exposure to military stressors could clarify whether theta-FRN is a consequence or predisposing factor for reexperiencing (Polusny et al., 2021). These data could also develop reduction of theta loss signaling as a biomarker for PTSD treatment response. For instance, if an individual’s reexperiencing symptoms were to improve, we would anticipate a corresponding reduction of their theta-FRN response to losses. This reduction would be expected to precede clinical symptom remission, reflecting a reduction in the salience of negative PEs over time. Additionally, value-based decision-making encompasses a range of processes beyond just valuation, such as prediction and action selection (Rangel et al., 2008). Future studies should capture neural activation during these other processes, possibly using gambling paradigms with semi-predictable outcomes like multi-armed bandits (O’Doherty et al., 2003) to gain deeper insight into associations with negative prediction biases. Finally, the predominance of males in our sample, reflecting the demographics of combat veterans seeking care at VA facilities, points to a need for future research to include more diverse samples, particularly with a higher representation of females who have well-characterized PTSD symptoms and drinking patterns.

In summary, our study shows mediofrontal theta elicited by losses exhibits opposing influences of intrusive reexperiencing and heavy drinking. This finding aligns with recent predictive coding models of PTSD (Kube et al., 2020; Putica et al., 2022), suggesting that chronic alcohol use might functionally reduce the intensity of salient negative prediction errors, thereby providing some relief from negative emotional reactivity. These insights not only deepen our understanding of the unique influences of PTSD and heavy drinking on brain salience signaling, but also suggest new avenues for neurobiologically-informed interventions. Specifically, treatments focusing on modulating mediofrontal theta activity (Chiang et al., 2022) could potentially address the exaggerated salience signaling associated with intrusive reexperiencing, offering a promising direction for future computationally-informed therapeutic approaches to PTSD management.

## Acknowledgments

This research was supported by the National Institutes of Health’s National Center for Advancing Translational Sciences, grants TL1R002493 and UL1TR002494 (ER), the Congressionally Directed Medical Research Program (Award number PT074550, contract W81XWH-08-2-0038 to SRS), and by the Department of Veterans Affairs Rehabilitation R&D Program, grant I01RX000622 (SRS). In addition, this project was supported with resources and the use of facilities at the Minneapolis VA Health Care System. The content is solely the responsibility of the authors and does not necessarily represent the official views or policy of the U.S. Department of Veterans Affairs, the United States Government, or the National Institutes of Health’s National Center for Advancing Translational Sciences.

## CRediT Author Statement

ER: Conceptualization, Methodology, Software, Validation, Formal Analysis, Investigation, Writing - Original Draft, Writing - Review and Editing, Visualization. CAM: Conceptualization, Methodology, Validation, Data Curation, Formal Analysis, Writing - Review and Editing. SF: Visualization, Validation. EB: Formal Analysis, Software, Validation, Methodology. SRS: Resources, Data Curation, Writing - Review and Editing, Supervision, Project Administration, Funding Acquisition.

